# An Intra-brainstem Circuit for Pain-induced Inhibition of Itch

**DOI:** 10.1101/2024.07.02.601724

**Authors:** Jagat Narayan Prajapati, Devanshi Piyush Shah, Arnab Barik

**Affiliations:** Center for Neuroscience, Indian Institute of Science, Bengaluru, India, 560012

## Abstract

Pain and itch are unpleasant and distinct sensations that give rise to behaviors such as reflexive withdrawal and scratching in humans and mice. Interestingly, it has been observed that pain modulate itch through the neural circuits housed in the brain and spinal cord. However, we are yet to fully understand the identities of and mechanisms by which specific neural circuits mediate pain-induced modulation of itch. Independent studies indicate that brainstem nuclei such as the lateral parabrachial nucleus (LPBN) and rostral ventromedial medulla (RVM) are important for the suppression of itch by painful stimuli. Here, using mouse and viral genetics, rabies tracing, chemogenetics, and calcium imaging, we show that the synaptic connections between LPBN and RVM plays an instrumental role in the interactions between pain and itch. Notably, we found that the LPBN neurons that express the gene encoding the substance P receptor, Tacr1 (LPBN^Tacr1^), synapse onto Tacr1-expressing RVM neurons (RVM^Tacr1^). The RVM^Tacr1^ neurons were found to be nociceptive, sufficient for inhibiting itch, and necessary for pain-induced itch suppression. Moreover, through brain-wide anterograde and retrograde viral tracing studies, we found that the RVM^Tacr1^ neurons are bidirectionally connected with LPBN, periaqueductal gray (PAG), and lateral hypothalamic area (LHA). Thus, together, our data indicate that the RVM^Tacr1^ neurons integrate nociceptive information to mediate itch-induced scratching and can mediate the physiological effects of itch through their downstream targets.

## Introduction

Scratching, which is mildly painful, suppresses itch, and prescription ointments containing capsaicin, the active chemical component of chili peppers, are effective in treating patients with chronic itch (Andersen et al., 2017; Mochizuki et al., 2017). Thus, the relationship between pain and itch is antagonistic. However, the neural substrate for this itch modulation remains unclear.

Previously, itch was considered to be a mild form of pain, and the underlying neural circuits that transmits and process pain and itch information were presumed to be the same (McMahon and Koltzenburg, 1992; Ikoma et al., 2006; Davidson and Giesler, 2010, Akiyama and Carstens, 2013). However, recent evidence points to the contrary. Distinct peripheral receptors are tuned to histamine-induced itch and are non-responsive to noxious mechanical or thermal stimuli (Schmelz et al., 1997). Notably, the ablation of a subpopulation of spinal cord interneurons impaired itch-induced scratching and not the behavioral responses to other high-threshold somatosensory stimuli (Sun and Chen, 2007; Sun et al., 2009). In contrast, convergence between the neural circuits, responsive to both pain and itch are observed in the brain. In the lateral parabrachial nucleus (LPBN), where the projection neurons from the spinal cord carrying nociceptive information terminate, neurons encode both pain and itch. LPBN neurons are primarily glutamatergic, and when the entire LPBN population is activated, it promotes itch, whereas silencing suppresses it (Mu et al., 2017). A subset of lateral amygdala projecting PBN neurons, expressing the neuropeptide Calcitonin gene-related peptide (CGRP), are activated by itch and, when silenced, abolished itch-induced scratching (Campos et al., 2018; Palmiter, 2018). Interestingly, LPBN neurons expressing the receptor for substance P, Tacr1 (LPBN^Tacr1^) are primarily nociceptive and yet, when stimulated, inhibit itch (Barik et al., 2021). The LPBN^Tacr1^ are synaptic targets of spinal cord Tacr1 expressing projection neurons that transmit noxious information to the brain (Deng et al., 2020; Barik et al., 2021). Thus, the spinal^Tacr1^-LPBN^Tacr1^ constitutes a circuitry that plays an important role in directing attention to painful stimuli if the animals are simultaneously exposed to itch stimuli. However, it is unclear how spinal^Tacr1^-LPBN^Tacr1^ circuitry inhibits itch through their downstream circuitries.

Recently, the rostral ventromedial medulla (RVM), a key nuclei for descending pain modulation, has been implicated in itch processing (Gao et al., 2019, 2021; Liu et al., 2019; Follansbee et al., 2022; Nguyen et al., 2022, 2023). Classical electrophysiological experiments have classified RVM neurons as ON, OFF, and NEUTRAL cells (Heinricher et al., 1989, 2009; De Preter and Heinricher, 2024), depending on their responses to noxious stimuli. Pruritogens activated and inhibited the pro-nociceptive ON and anti-nociceptive OFF cells, respectively (Follansbee 2018). Further, scratching similarly activated and inhibited the ON and OFF cells, indicating the involvement of RVM neurons in itch modulation (Follansbee 2018). Thus, a high-threshold noxious mechanical stimuli such as scratching can suppress itch through the RVM neurons. Interestingly, recent studies have shown that Tacr1 is expressed in the RVM, and the activation of RVM^Tacr1^ inhibited itch (Follansbee et al., 2022). We hypothesized that the LPBN^Tacr1^ neurons synapse onto RVM^Tacr1^ neurons to suppress itch. To that end, we found that the LPBN^Tacr1^ neurons have robust projections to the RVM, the LPBN^Tacr1^, and RVM^Tacr1^ neurons are mono-synaptically connected, and the RVM^Tacr1^ neurons, as in the LPBN^Tacr1^ neurons, when activated, inhibit itch. Notably, we found that the RVM^Tacr1^ neurons are necessary for pain-induced suppression of itch. Thus, a molecularly defined intra-brainstem circuit between the LPBN and the RVM plays a pivotal role in pain modulation of itch.

## Results

### Activation of genetically defined cell types in PBN inhibits itch

We previously found that the LPBN^Tacr1^ neurons are sufficient for the promotion of nocifensive behaviors and suppression of itch (Barik et al., 2021). The spinal neurons expressing the gene encoding the neuropeptide Gastrin-releasing peptide (Grp) receptor (Grpr) are critical for itch transmission (Sun and Chen, 2007; Sun et al., 2009). It was not known if the Grpr gene is expressed in the LPBN, and whether the neurons which express Grpr play any role in itch modulation. Hence, we performed multiplex fluorescent *in situ* hybridization (RNAscope) to visualize the Grpr and Tacr1 expressing neurons in PBN (Fig. 1A). We observed the expression of *grpr* and *tacr1* mRNA in the external-lateral part of the LPBN (Fig. 1A top center and right). *grpr* and *tacr1* expressing neurons partially overlapped (Fig. 1A bottom middle). We found that both the neuronal populations are glutamatergic, and thus they express the excitatory neuronal marker *slc32a*, the gene encoding the vesicular glutamate transporter (VGlut2) (Barik et al., 2018; Pauli et al., 2022). Next, we chemogenetically activated the LPBN Grpr (LPBN^Grpr^) expressing neurons to test if the manipulation of these neurons phenocopy the activation of LPBN^Tacr1^ neurons. To that end, we injected the Adeno-associated virus (AAV) encoding Cre-dependent hM3Dq in the LPBN of Grpr^iCre^ mice (Alexander et al., 2009; Krashes et al., 2011) to express the activating chemogenetic Designer Receptors Exclusively Activated by Designer Drugs (DREADDs) hM3Dq specifically in the PBN^Grpr^ neurons (Fig. 1B&C). hM3Dq expression in the PBN^Grpr^ neurons was confirmed by the expression of mCherry, which was fused with the hM3Dq protein (Fig. 1C). As predicted, activation of the PBN^Grpr^ neurons (Nagai et al., 2020) (Fig. 1D) suppressed chloroquine-induced scratching behavior in mice, similar to when PBN^Tacr1^ neurons were stimulated. This was expected as Grpr and Tacr1 are expressed in overlapping neuronal population in the LPBN. Hence, our data indicate that activating two genetically defined cell types (PBN^Grpr^ and PBN^Tacr1^) in the LPBN is sufficient to suppress itch and may play an important role in pain-induced itch modulation.

**Fig. 1.**
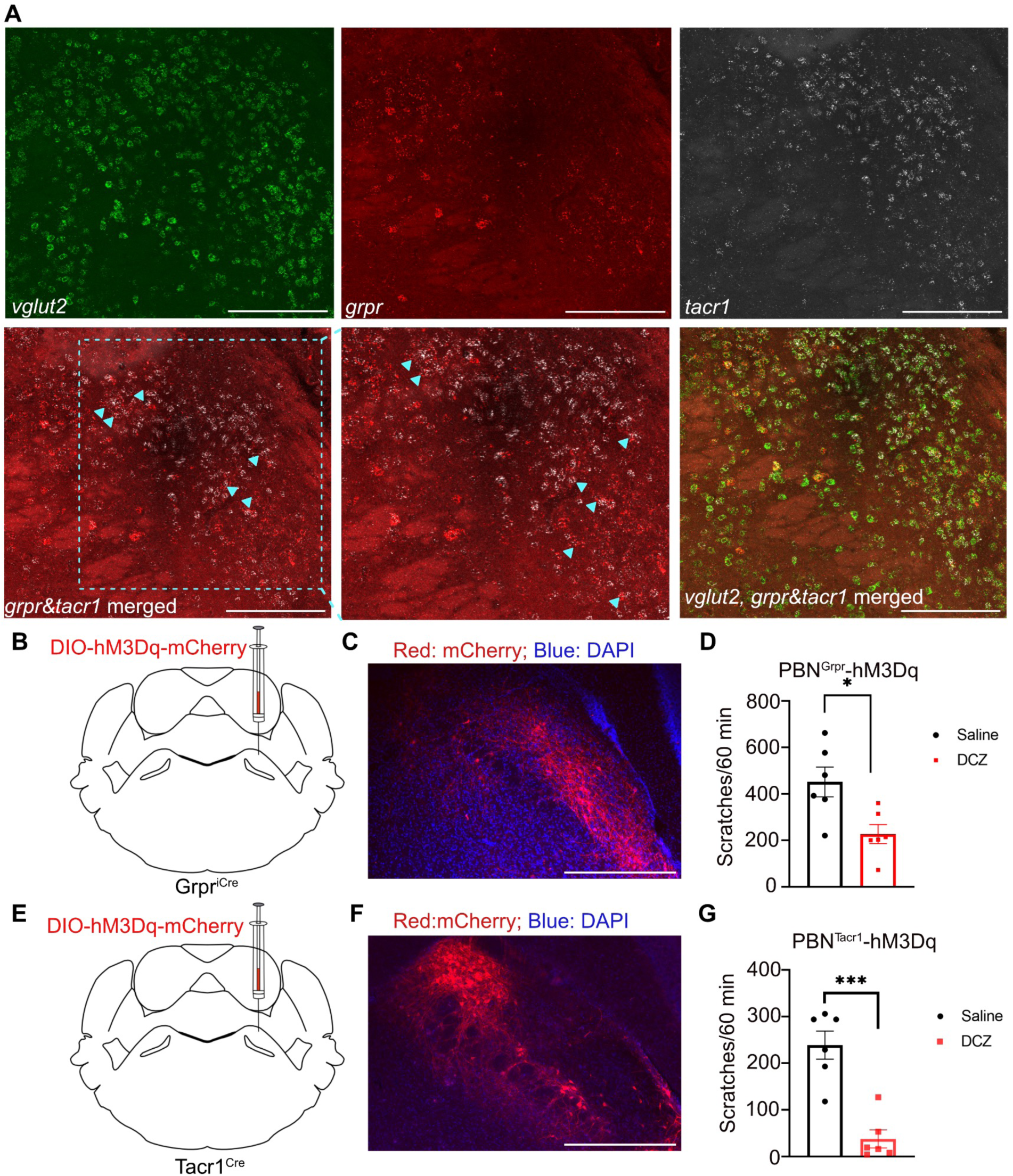
Activation of genetically defined cell types in PBN inhibits itch. (A) Multiplexed fluorescent *in situ* hybridization for *grpr* (red), *tacr1* (grey), and *slc17a6* (green) mRNA in the LPBN, scale bar 100 um. Magenta arrows mark the overlap between *grpr* and *tacr1*. (B) Injection of AAV expressing Cre-dependent excitatory DREADD hM3Dq into the LPBN of Grpr^iCre^ mice. (C) Expression of hM3Dq in PBN^Grpr^ neurons (mCherry-red, DAPI-blue), scale bar 500 µm. (D) Effect of i.p. DCZ (t-test, 224.2 ± 65.30, \**P*=0.0186, n=6) administration on chloroquine-induced scratching compared with i.p. saline. (E) Injection of AAV expressing Cre-dependent excitatory DREADD hM3Dq into the LPBN of Tacr1^Cre^ mice. (F) Expression of hM3Dq in PBN^Tacr1^ neurons (mCherry-red, DAPI-blue), scale bar 500 µm. (G) Effect of i.p. DCZ (t-test, 201.2 ±27.03 \*\*\**P*=0.0007, n=6) administration on chloroquine-induced scratching compared with i.p. saline.

### PBN^Grpr^ and PBN^Tacr1^ neurons project to the RVM

The diverse output of the PBN to different brain regions drives physiological and behavioral responses to pain and thus helps in maintaining homeostasis and promoting survival (Barik et al., 2018; Campos et al., 2018; Palmiter, 2018; Chiang et al., 2019, 2020; Deng et al., 2020). To delineate the mechanisms of suppression of itch by LPBN^Grpr^ and LPBN^Tacr1^ activation, we mapped out the brain-wide projections of LPBN^Grpr^ and LPBN^Tacr1^ neurons (Fig. 2). We used the AAV-mediated anterograde tracing technique and expressed the cell-filling fluorescent protein tdTomato in the PBN^Grpr^ and PBN^Tacr1^ neurons (Fig. 2A&M). We found robust tdTomato expression in the LPBN of both the strains (Fig. 2 B&C and Fig. 2N&O). Serial sectioning and sequential whole-brain imaging of the brain tissues expressing tdTomato in the LPBN^Grpr^ neurons and LPBN^Tacr1^ neurons showed axon terminals in various brain areas implicated in pain processing and nocifensive behaviors. LPBN^Grpr^ and LPBN^Tacr1^ neurons both projected to the lateral preoptic area (LPO), anterior hypothalamic area (AHC), amygdala, and lateral hypothalamus (LH) in the forebrain (Fig. 2F-I and Fig.2 R-U). In the brainstem, we observed projections in the PAG (periaqueductal gray), substantia nigra (SNR), the nucleus of the solitary tract, parvocellular reticular nucleus (PCRt), and the medullary reticular nucleus (MdD) (Fig. 2J-L, and Fig. 2V-X). Importantly, we found that the LPBN^Grpr^ and LPBN^Tacr1^ neurons both project to the RVM (rostral ventromedial medulla) (Fig. 2D, E, P, Q). Thus, we observed similar brain-wide projections of LPBN^Grpr^ and LPBN^Tacr1^ neurons. This finding is in agreement with the overlapping nature of *Tacr1* and *Grpr* expression in the LPBN (Fig. 1A). However, we observed stronger projections of LPBN^Grpr^ neurons compared to the LPBN^Tacr1^ neurons at the target regions, possibly due to the higher number of *Grpr* expressing neurons in the LPBN.

**Fig. 2.**
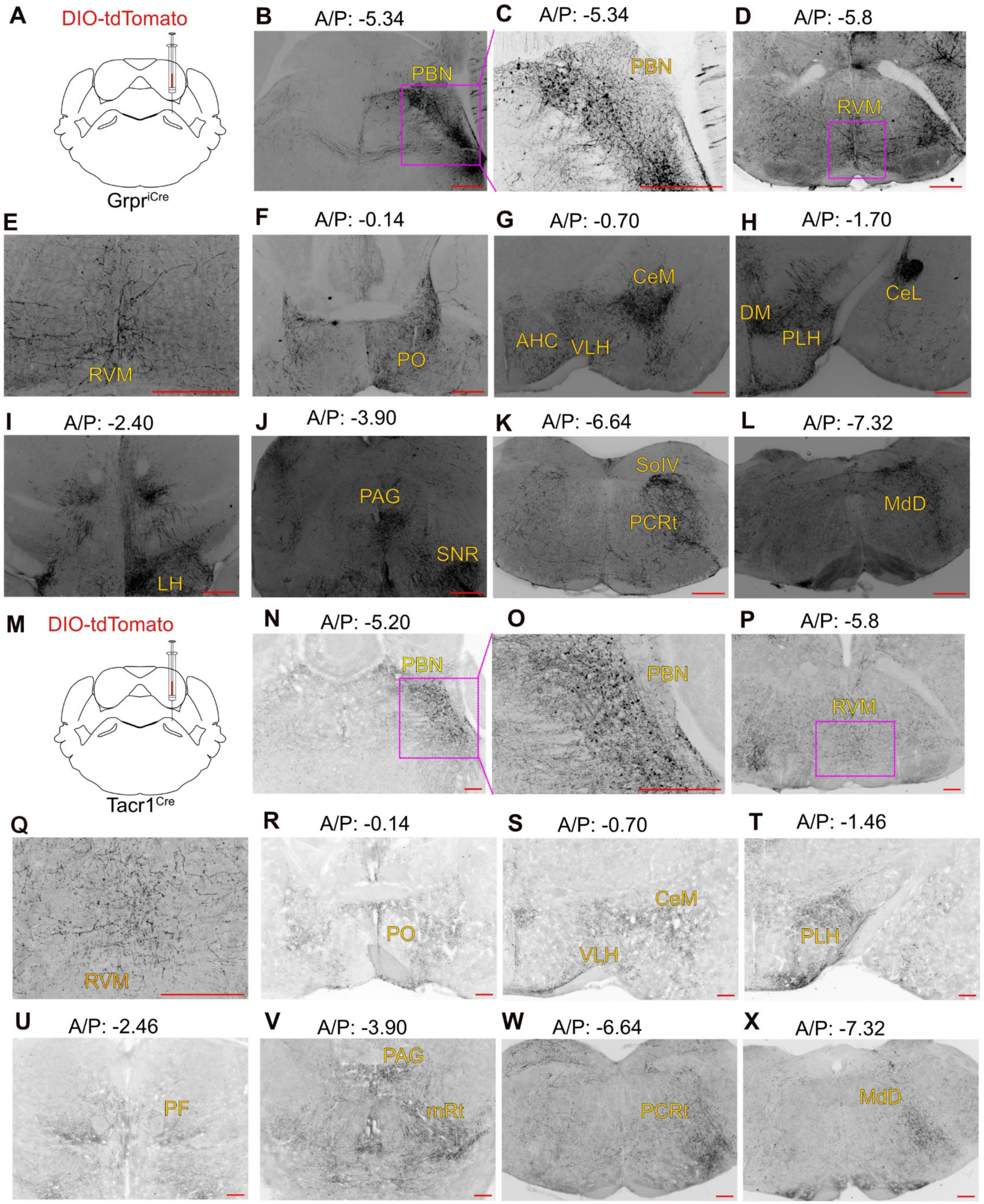
PBN^GRPR^ and PBN^Tacr1^ neurons project to RVM. (A) AAV encoding Cre-dependent tdTomato was stereotaxically injected into the PBN of Grpr^iCre^ mice. (B) Expression of tdTomato in PBN^Grpr^ neurons. (C) Higher magnification image of PBN^Grpr^ neurons. (D-L) PBN^Grpr^ neuron projections across various brain regions. (M) AAV encoding Cre-dependent tdTomato was stereotaxically injected into the PBN of Tacr1^Cre^ mice. (N) Expression of tdTomato in PBN^Tacr1^ neurons. (O) Higher magnification image of PBN^Tacr1^ neurons. (P-X) PBN^Tacr1^ neuron projections across various brain regions, scale bar 500 µm.

### PBN^Tacr1^ neurons are synaptically connected to the RVM^Tacr1^ neurons

As mentioned before, targeted activation of the Tacr1-expressing neurons in the RVM (RVM^Tacr1^) suppressed non-histaminergic itch in mice (Follansbee et al., 2022). Moreover, nociceptive inputs to the RVM were shown to be derived from the LPBN (Roeder et al., 2016; Chen et al., 2017). Since we found that the LPBN^Tacr1^ neurons project to the RVM, we hypothesized that the LPBN^Tacr1^ neurons may synapse with the RVM^Tacr1^ neurons. To test the synaptic connectivity between these two groups of neurons, we injected AAVs encoding Cre-dependent synaptophysin-tagged Green fluorescence protein (DIO-SynGFP) in the LPBN and DIO-tdTomato in the RVM (Fig. 3A). This approach labelled the synaptic vesicles of the LPBN neurons with GFP, as the GFP is fused with the synaptophysin protein that is specifically enriched in the synaptic vesicles in the axon terminals (Wiedenmann and Franke, 1985; Barik et al., 2018) (Fig. 3A) of the neurons that are to be labelled. In the same mice, the RVM^Tacr1^ neurons were labelled with cell-filling tdTomato fluorescent protein (Fig. 3A). Syn-GFP and tdTomato were closely apposed in the RVM, indicating putative synapses between the PBN^Tacr1^ and RVM^Tacr1^ neurons (Fig. 3C&D). To further consolidate our findings, we injected AAVs encoding DIO-SynGFP in the LPBN and DIO-PSD95-tagRFP in the RVM of Tacr1-Cre mice. PSD95 is a protein, that is enriched in the post-synaptic density of excitatory synapses and, when tagged with fluorescent proteins, can be useful in visualizing synapses in vivo (Barik et al., 2021; Quillet et al., 2023). Upon imaging the synapses on the RVM^Tacr1^ neurons with traditional confocal and super-resolution microscopy, we found that the PSD95-tagRFP and Syn-GFP were closely apposed (Fig. 3G&H), indicating that synaptic connections exist between the RVM and LPBN Tacr1 expressing neurons. Thus, using complementary genetic anatomical strategies, we demonstrate that the RVM^Tacr1^ neurons are downstream of the LPBN^Tacr1^ neurons.

**Fig. 3.**
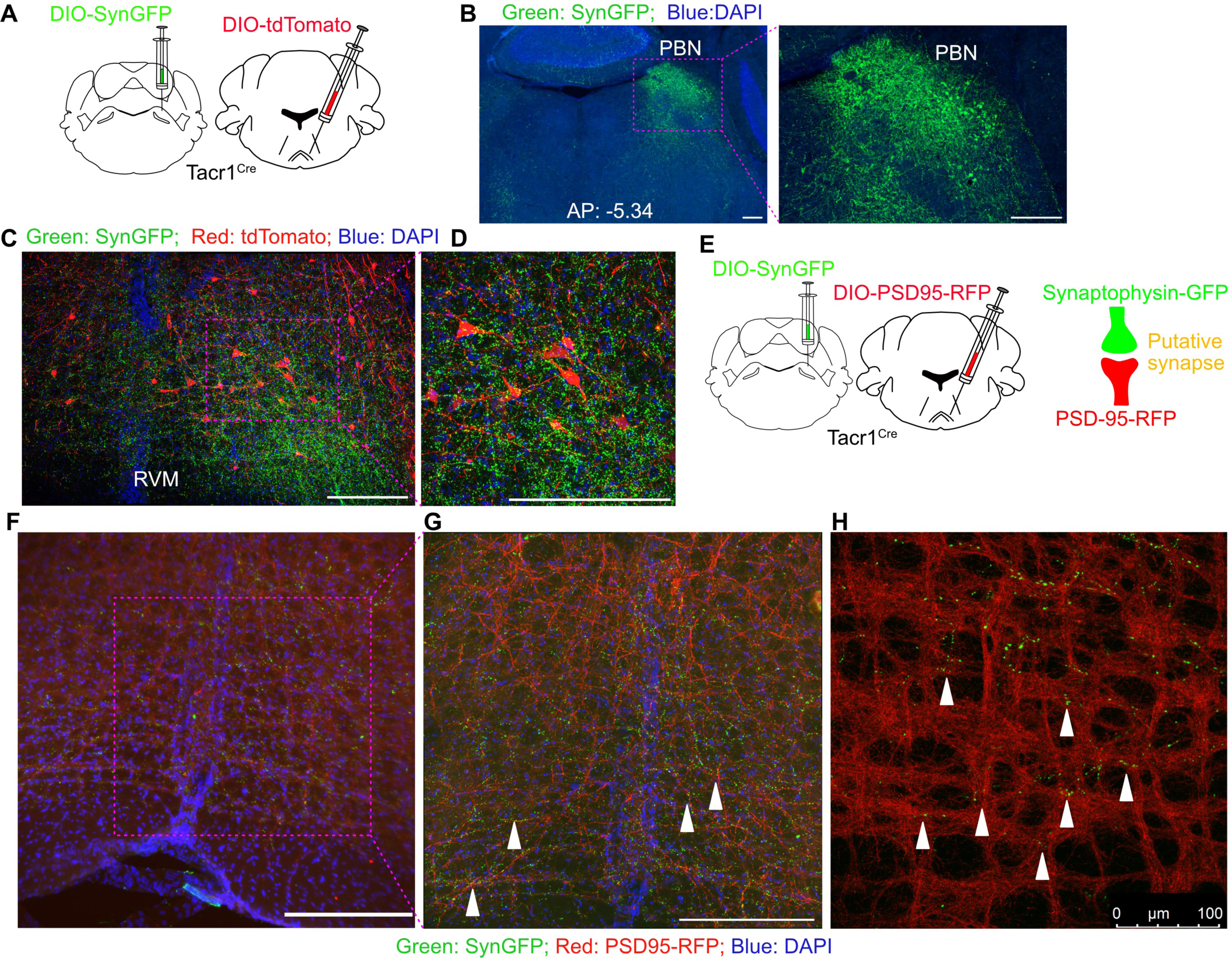
PBN^Tacr1^ neurons are synaptically connected to the RVM^Tacr1^ neurons. (A) AAV expressing Cre-dependent synaptophysin tagged GFP (SynGFP) and Cre-dependent tdTomato were injected into the LPBN and RVM of Tacr1^Cre^ mice, respectively. (B) Expression of SynGFP in PBN^Tacr1^ neurons (SynGFP-green, DAPI-blue). (C) Expression of tdTomato in RVM^Tacr1^ neurons (tdTomato-red, SynGFP-green, DAPI-blue). (D) High magnification representation of RVM^tdtomato+^ neurons. (E) AAV encoding Cre-dependent SynGFP was injected into the PBN, and AAV encoding PSD-95 tagRFP was injected into the RVM of Tacr1^Cre^ mice. (F) Schematic showing overlap between SynGFP and PSD-95 RFP marks putative synapse. (G&H) Overlap seen between green and red puncta (marked with white arrows) in the RVM (SynGFP-green, PSD-95 tagRFP-red, DAPI-blue), G, 40X Confocal; H, 40X AiryScan, scale bar 250 µm.

### RVM^Tacr1^ neurons suppress acute itch and sustained thermal pain without affecting the reflexive withdrawal thresholds

Next, we chemogenetically stimulated the RVM^Tacr1^ neurons to observe its effect on pain and itch behaviors. To that end, we injected AAV encoding Cre-dependent hM3Dq into the RVM of Tacr1^Cre^ mice (Fig. 4A). Three weeks later, robust expression of mCherry was found in the RVM, suggesting successful labelling of the RVM^Tacr1^ neurons with hM3Dq (Fig. 4B). DCZ (deschloroclozapine) was administered i.p. to chemogenetically stimulate the RVM^Tacr1^ neuron. Since the RVM^Tacr1^ neurons are downstream of the LPBN^Tacr1^ neurons (Fig. 3), and targeted activation of LPBN^Tacr1^ neurons suppresses chloroquine-mediated itch (Fig. 1), we expected chemogenetic activation of RVM^Tacr1^ neurons to be sufficient for modulating itch. Indeed, the acute itch was suppressed in DCZ and not in the saline-administered mice expressing hM3Dq in the RVM^Tacr1^ neurons (Fig. 4C). In addition, since RVM is traditionally known to be involved in descending pain modulation, we tested the role of RVM^Tacr1^ neurons in behavioral responses to noxious thermal stimuli. We subjected the mice to the hot-plate test (Espejo and Mir, 1993; Reddy et al., 2023) at 52°C for one minute, and evaluated their latency and frequency of the paw licks, shakes, and jumps. Activation of RVM^Tacr1^ neurons caused a marked reduction in the number of licks, shakes, and jumps in mice, suggesting an analgesic effect on sustained thermal pain (Fig. 4D, F&H). Similarly, the activation of RVM^Tacr1^ neurons increased the latency to lick (Fig. 4E) and shake (Fig. 4G). In contrast, we found that the latency to jump was not affected by the activation of RVM^Tacr1^ neurons (Fig. 4I). Notably, we found that the activation of RVM^Tacr1^ neurons fails to affect the tail flick latency (Fig. 4 J), suggesting that these RVM^Tacr1^ neurons may not play an important role in determining the reflexive nociceptive thresholds. Together, our data indicate that the RVM^Tacr1^ neurons, when analgesic, can suppress the supraspinal behavioral responses to noxious thermal stimuli without affecting the spinal reflexes.

**Fig. 4.**
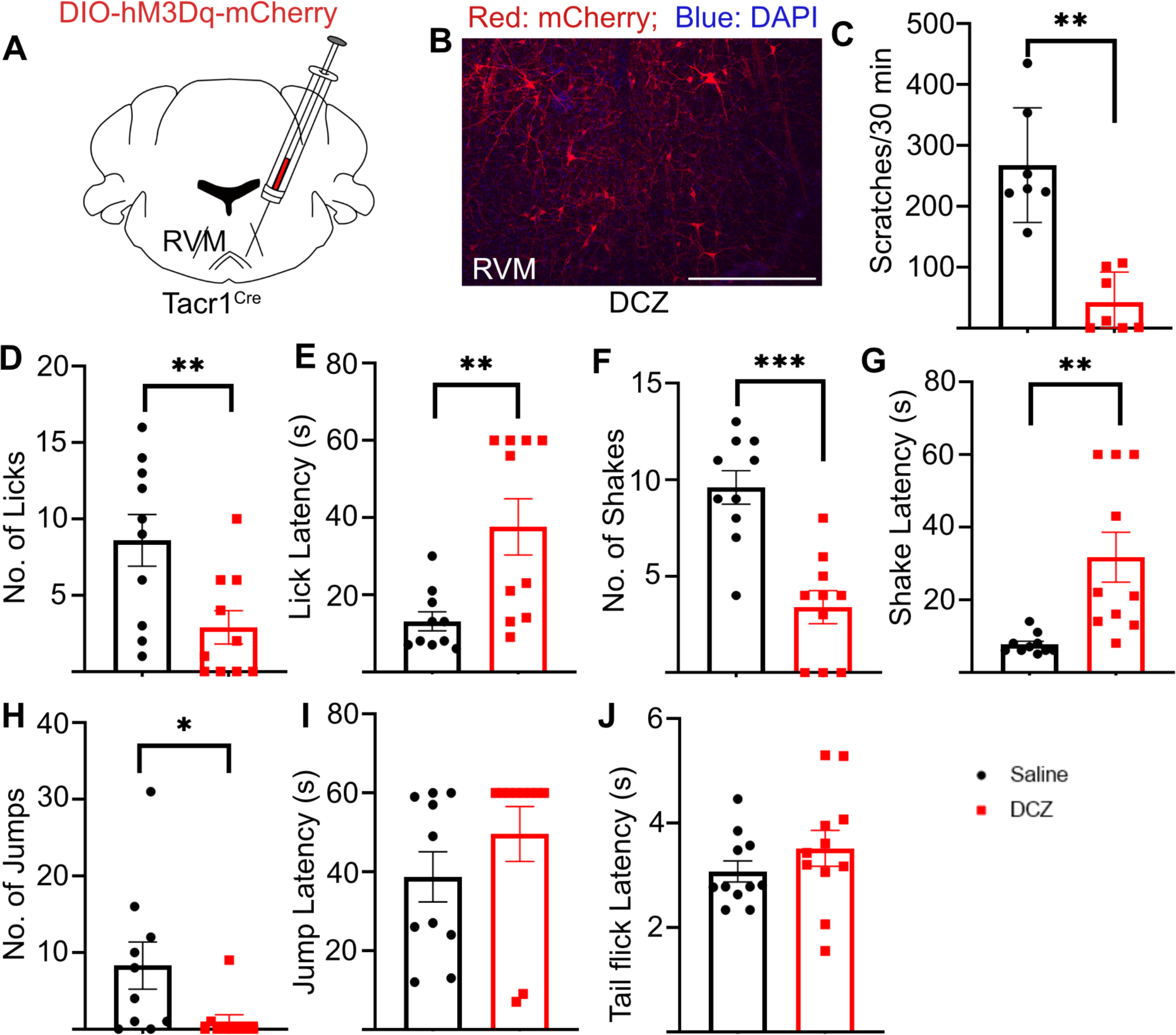
Activation of RVM^Tacr1^ neurons suppresses acute itch and sustained pain. (A) RVM^Tacr1^ neurons were labelled with excitatory DREADD hM3Dq in Tacr1^Cre^ mice. (B) Expression of hM3Dq in RVM^Tacr1^ neurons (mCherry-red, DAPI-blue). (C) Effect of i.p. DCZ on the no. of scratches (t-test, 225.6 ± 37.87, \*\**P*=0.001, n=7) compared with i.p. saline. (D-I) Effect of i.p. DCZ administration on the no. of licks (t-test, 5.7±1.2, \*\**P*=0.0011, n=10), lick latency (t-test, 24.50 ± 5.928, \*\**P*=0.0025, n=10), no. of shakes (t-test, 6.2 ± 1.104, \*\*\**P*=0.0003, n=10), shake latency (t-test, 24 ± 6.896, \*\**P*=0.0069, n=10), no. of jumps (t-test, 7.3 ± 3.138, \**P*=0.045, n=10), jump latency, and (J) tail flick latency (n=11) compared with i.p. saline.

### RVM^Tacr1^ neurons are necessary for pain-induced inhibition of itch

Since we found that the RVM^Tacr1^ neurons were sufficient to suppress itch (Fig. 4), we next tested if they are RVM^Tacr1^ neurons required for pain-mediated suppression of itch. To that end, we expressed the inhibitory chemogenetic DREADD hM4Di in the RVM^Tacr1^ neurons (Fig. 5A) and confirmed its expression by observing mCherry (fused with the hM4Di) in these neurons in the RVM (Figure 5C). To test the effect of pain on itch in these mice, we performed intraplantar injections of 0.5% (w/v) AITC (Allyl isothiocyanate). AITC is a known agonist of the TRPA1 channel and induces mild pain in mice and humans (Jordt et al., 2004; Bautista et al., 2006). We found that in the mice expressing hM4Di in the RVM^Tacr1^ neurons, under control conditions, i.e. when saline was administered i.p., intra-plantar AITC suppressed chloroquine-induced itch (Fig. 4D). Meanwhile, DCZ abrogated the AITC-mediated itch suppression in the same animals (Fig. 4D). Thus, from these experiments, we concluded that the RVM^Tacr1^ neurons play a facilitatory role and are required for pain-induced itch modulation.

**Fig. 5.**
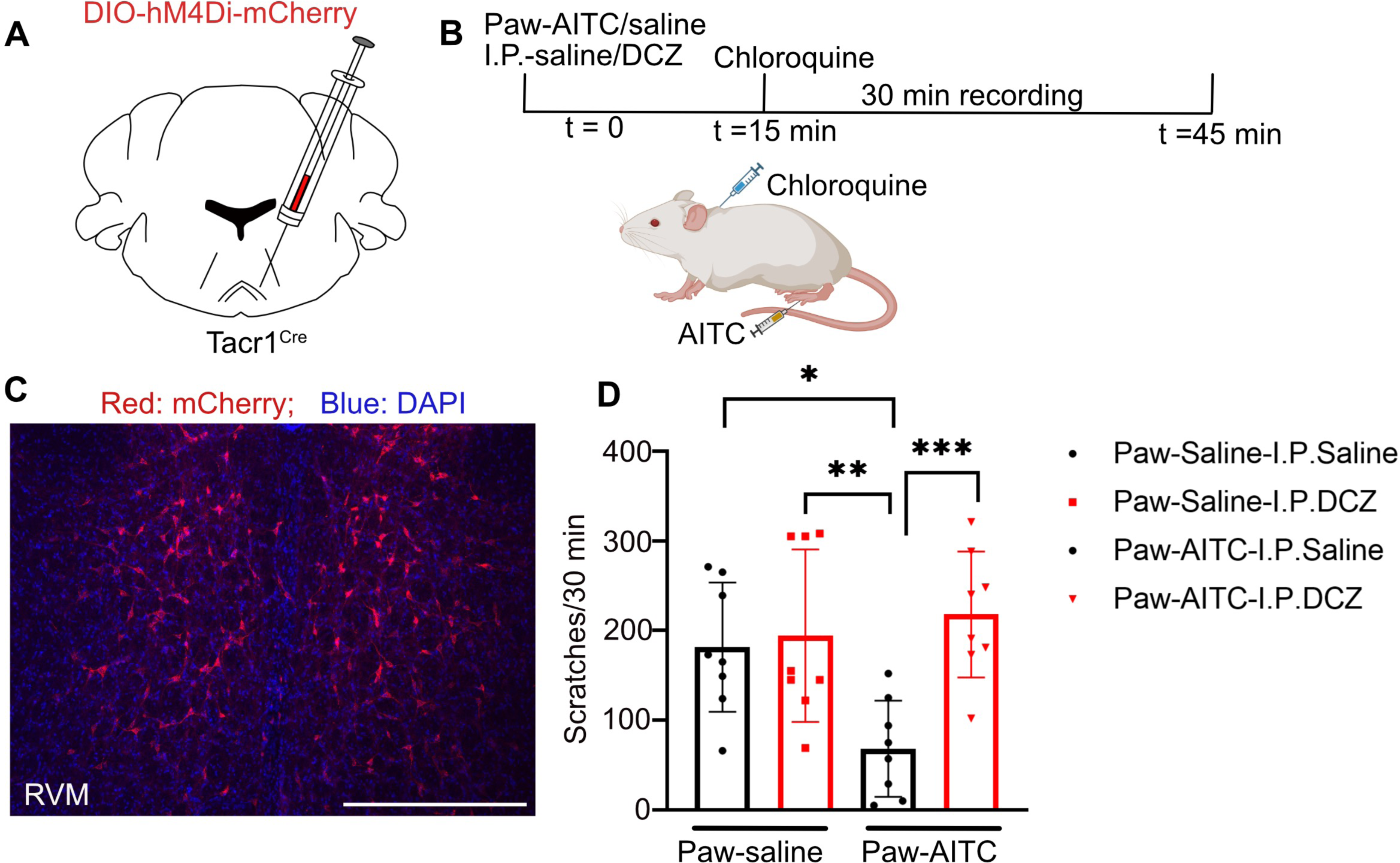
Inhibition of RVM^Tacr1^ neurons abolishes pain-induced inhibition of itch. (A) AAV encoding inhibitory DREADD hM4Di was injected in the RVM of Tacr1^Cre^ mice. (B) Schematic representation of experiment timeline and order. (C) Expression of hM4Di in RVM^Tacr1^ neurons (mCherry-red, DAPI-blue). (D) Inhibition of RVM^Tacr1^ neurons abolishes AITC-induced inhibition of itch (Two-way ANOVA test, multiple comparison Bonferroni, paw-saline-i.p.-saline vs. paw-AITC-i.p.-saline-113.1 ± 33.73, *P*=0.018; paw-saline-i.p.-DCZ vs. paw-AITC-i.p.-saline-125.9 ± 33.73, *P*=0.0074; paw-AITC-i.p.-saline vs. paw-AITC-i.p.-DCZ-149.6 ± 33.73, *P*=0.0014; n=8).

### RVM^Tacr1^ neurons are tuned to noxious thermal pain

In the next set of experiments, we sought to determine the response profiles of the RVM^Tacr1^ neurons when exposed to noxious stimuli. To that end, we recorded the activity of RVM^Tacr1^ neurons while the mice were subjected to the hot-plate test using fiber photometry (Gunaydin et al., 2014). AAVs encoding Cre-dependent GCaMP6s were stereotaxically infused into the RVM of Tacr1^Cre^ mice (Fig. 6A). An optical fiber cannula was implanted just above the RVM three weeks after the viral infusion to record the GCaMP6s fluorescence. Robust expression of GCaMP6s was found in the RVM^Tacr1^ neurons (Fig. 6B). We found that increased activity in the RVM^Tacr1^ neurons coincided with licking and shaking of the paw in response to the noxious heat. When we recorded RVM^Tacr1^ neural activity on the hot plate, set in a ramp configuration, where the plate temperature increased from 32°C-56°C over 5 minutes, we found that the activity in the RVM^Tacr1^ neurons increased when the temperature reached the unbearable noxious range (Caterina et al., 1997, 2000) (Fig. 6F&G). On the ramped hot plate test, above 52°C, we found that mice jump in efforts to escape the enclosure (Fig. 6C, 4H). However, we did not observe a correlation between RVM^Tacr1^ activity and the escape responses (Fig. 6D-G). In sum, we found that the RVM^Tacr1^ neurons are tuned to somatosensory stimuli in the noxious range, and the activity coincided with coping behaviors such as licks and shakes.

**Fig. 6.**
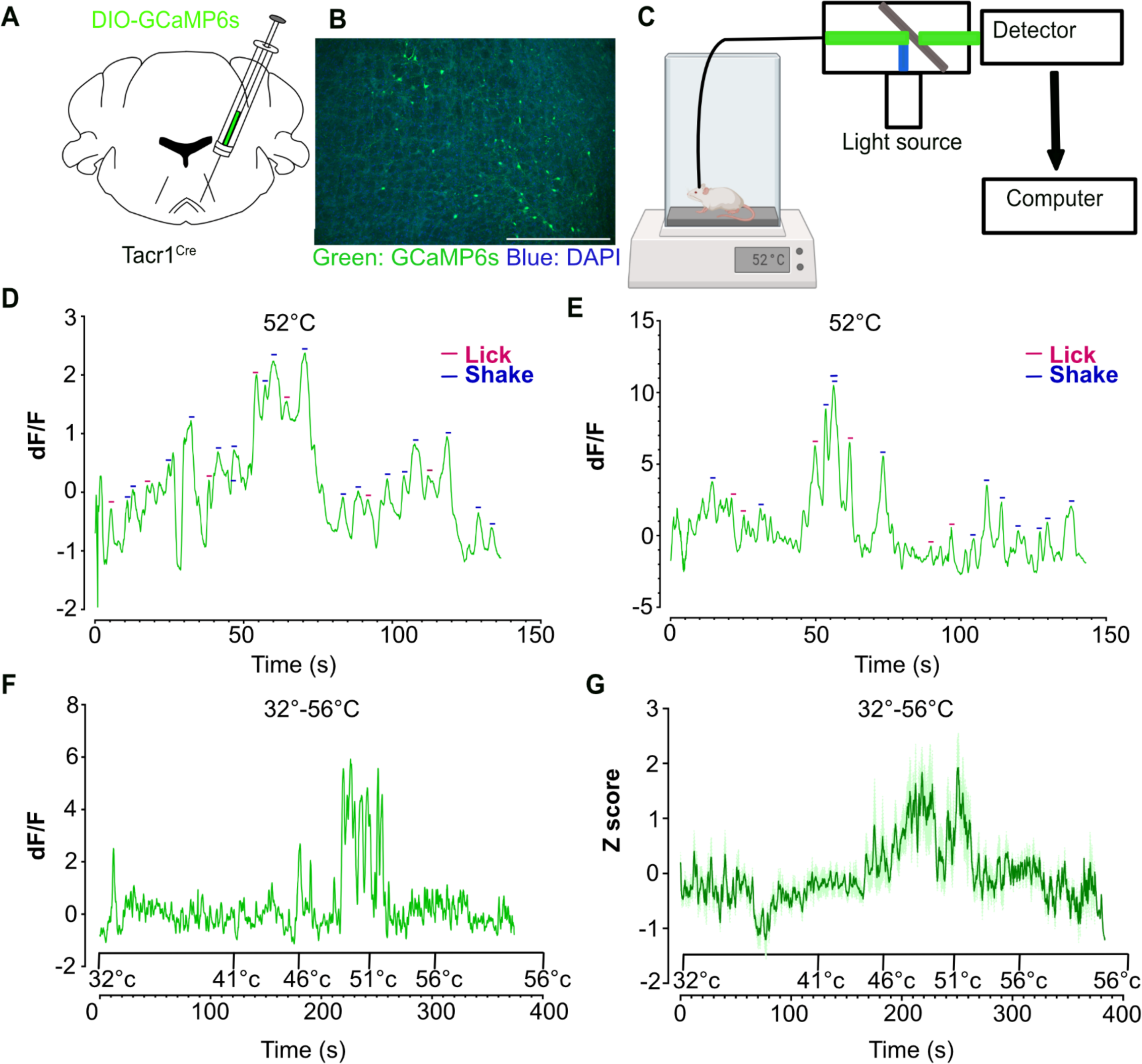
Activation of RVM^Tacr1^ neurons suppresses acute itch and sustained pain. (A) AAV encoding Cre-dependent GCaMP6s was injected into the RVM of Tacr1^Cre^ mice. (B) Expression of GCaMP6s (green-GCaMP6s, DAPI-blue) in RVM^Tacr1^ neurons. (C) Schematic showing the calcium activity recording from RVM^Tacr1^ neurons while mice were challenged on a hotplate test. (D&E) Representative traces from two mice challenged on the hotplate at 52°C (n=2). The peaks correspond to nocifensive behavior licks and shakes (red-licks, blue-shakes). (F) Representative trace of calcium activity when mice were challenged to 32°-56°C gradient on the hotplate (n=1). (G) Z score of neural activity when mice were challenged to 32°-56°C gradient on the hotplate (n=4).

### Brain-wide mapping of RVM^Tacr1^ inputs and outputs

RVM is known to project to the superficial as well as deeper spinal laminas via the dorsolateral and ventrolateral funiculus to modulate nociceptive transmission (Fields et al., 1995; François et al., 2017; Nguyen et al., 2022; Otsu and Aubrey, 2022; Ganley et al., 2023). However, the ascending projections of RVM to the rest of the brain, in the brainstem, midbrain, and forebrain, remain poorly explored. We injected AAVs carrying gene sequences for Cre-dependent cell-filling GFP into the RVM of Tacr1^Cre^ mice (Fig. 7A). We observed robust expression of GFP in these neurons after three weeks of viral infection (Fig. 7B). The GFP expressing axon terminals of the RVM^Tacr1^ neurons were observed in the medial septum (MS), nucleus of the vertical limb of the diagonal band (VDB), medial preoptic area (MPA), and peduncular part lateral hypothalamus (PLH) (Fig. 7C-E), in the forebrain. In the diencephalon, RVM^Tacr1^ neurons projected to the central medial thalamus (CT), parafascicular thalamus (PF), and lateral hypothalamus (LHA) (Fig. 7F-H). The RVM^Tacr1^ neurons were found to project to the PAG, RF, and LPBN in the brainstem (Fig. 7I-K). In addition to the mentioned brain regions, the RVM^Tacr1^ neurons projected to the superficial lamina in the dorsal horn of the spinal cord in the cervical and lumbar regions (Fig. 7L&M). We observed that most of the GFP labeled terminal of RVM^Tacr1^ neurons were dorsal to the IB4 stained (red, marked by white arrows in Fig. 7L&M) regions in the spinal cord, suggesting that RVM^Tacr1^ neurons project to lamina I of the dorsal horn. Furthermore, the RVM^Tacr1^ projections were also found in spinal lamina V, the ventral horn (more prominent in the cervical region than the lumbar), and the lamina X surrounding the central canal of the spinal cord (Fig. 7L&M). Thus, the AITC-mediated pain suppression, facilitated by the RVM^Tacr1^ neurons, might be through their synaptic targets across the brain and the spinal cord.

**Fig. 7.**
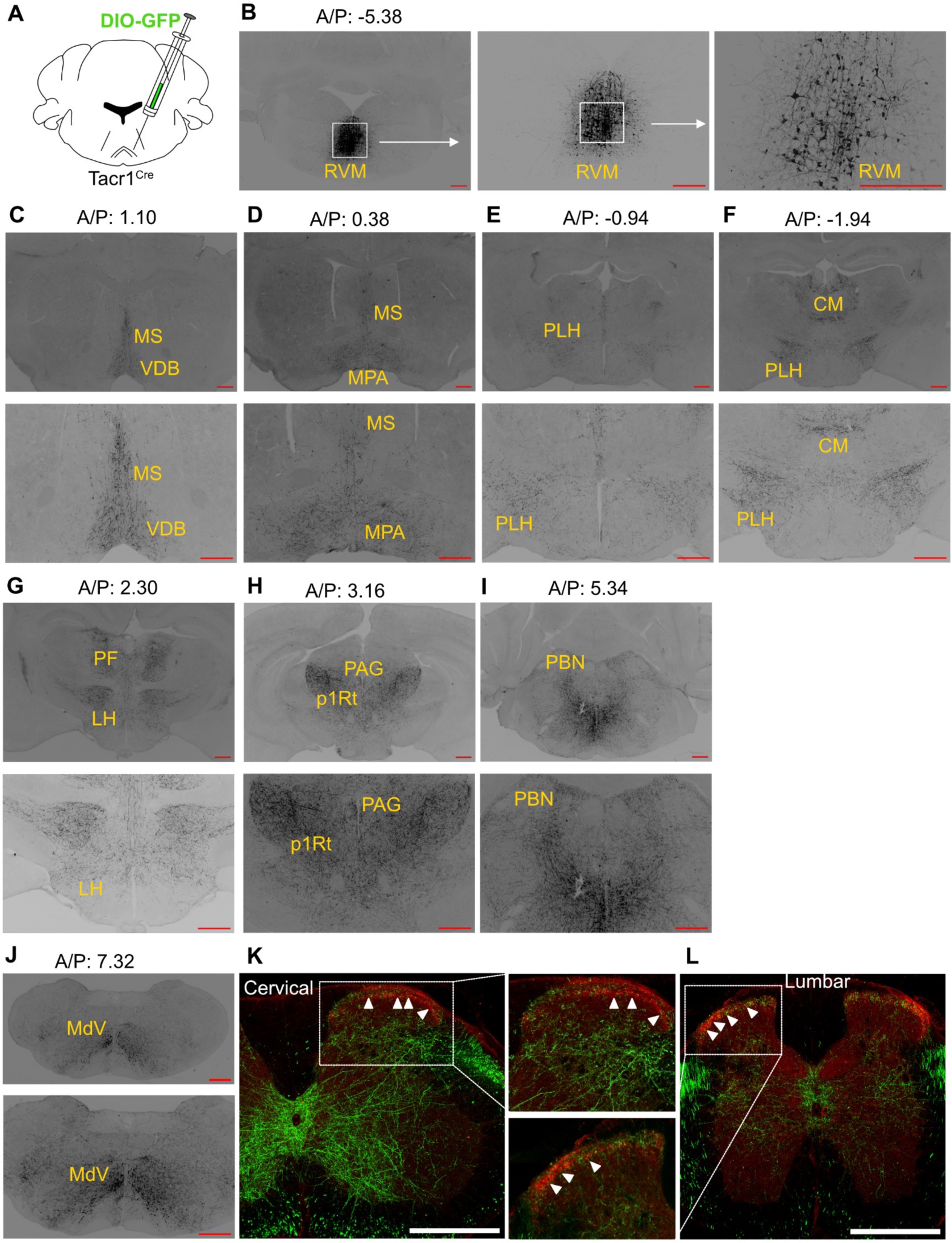
Projections of RVM^Tacr1^ neurons. (A) AAV encoding Cre-dependent GFP was injected into the RVM of Tacr1^Cre^ mice. (B) Expression of GFP in RVM^Tacr1^ neurons (black cell bodies). Subsequent images are higher magnification representations of the same RVM neurons, scale bar 500 µm. (C-K) Brain-wide projections of RVM^Tacr1^ neurons in 2X (top row) and 4X (bottom row) magnifications, scale bar 500µm. (L&M) Projections of RVM^Tacr1^ neurons to the dorsal horn of the spinal cord in cervical and lumbar regions, respectively. IB4 staining (red, indicated by white arrows), scale bar 500 µm.

Next, we sought to map the inputs to the RVM^Tacr1^ neurons. We used the monosynaptic rabies tracing technique (Callaway and Luo, 2015), which enables cell-type-specific labeling of direct synaptic inputs. Two AAVs, one expressing the rabies virus glycoprotein G and the other expressing the TVA receptor tagged with GFP in a Cre-dependent fashion, were stereotaxically injected into the RVM of Tacr1^Cre^ mice (Fig. 8A). Next, we infused the modified and pseudotyped rabies virus (expressing EnvA, deleted glycoprotein G and red fluorescent protein dsRed) into the RVM of the same mice three weeks later, in order to allow optimal expression of glycoprotein G and the TVA receptor (Fig. 8A). The starter RVM^Tacr1^ neurons expressed both GFP and dsRed (yellow) (Fig. 8B), as the neurons get infected by the rabies virus using TVA-EnvA (receptor-mediated endocytosis). The retrogradely labelled neurons expressed dsRed. We found that the RVM^Tacr1^ neurons receive inputs from the lateral hypothalamus (LH), ventral tegmental area (VTA), PAG, Superior Colliculus (SC), midbrain reticular formation (MRt), PBN, and the intermediate reticular nucleus (IRt) (Fig. 8C-I). Thus, the RVM^Tacr1^ neurons integrate nociceptive and physiological information from the brainstem and midbrain structures to modulate nocifensive responses to pain and itch.

**Fig. 8.**
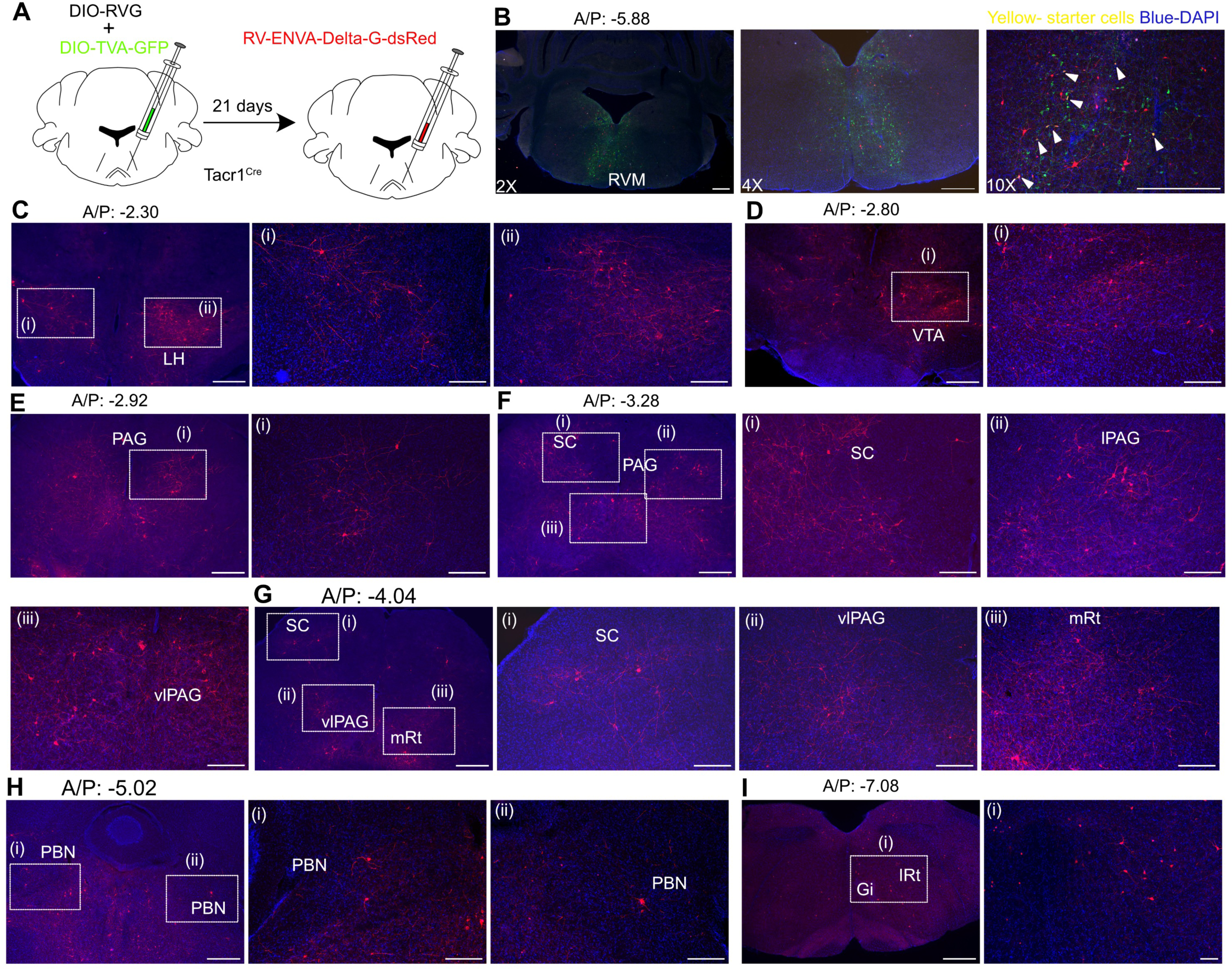
Presynaptic inputs of RVM^Tacr1^ neurons. (A) Experimental approach for monosynaptic rabies tracing. AAV expressing Cre-dependent TVA-GFP, and rabies glycoprotein G was injected into the RVM of Tacr1^Cre^ mice. 3 weeks later modified rabies virus with deleted glycoprotein G was injected into the RVM of the same Tacr1^Cre^ mice. (B) Site of injection (TVA-green, DAPI-blue). Starter cells (yellow, indicated by white arrows) express both GFP and dsRed. (C-I) Monosynaptic input of RVM^Tacr1^ neurons from diverse brain regions, scale bar 500 µm.

## Discussion

In this study, we delineated a novel neural circuitry in the brainstem that mediates pain modulation of itch. We found that Tacr1-expressing neurons in the LPBN, which receive noxious somatosensory information from the spinal cord and can cause pain, are sufficient to suppress itch. This itch modulation is carried out through their postsynaptic targets in the RVM, which also expresses Tacr1. The RVM^Tacr1^ neurons were found to be sufficient in suppressing itch and necessary for AITC-induced pain modulation of itch. Further, we found that the RVM^Tacr1^ neurons were tuned to noxious thermal stimuli and the coping responses due to the pain. Further, we anatomically mapped the inputs and outputs of these neurons. We found that the RVM^Tacr1^ neurons are bidirectionally connected to brain areas implicated in pain processing and determining brain states related to stress and anxiety. Together, we have delineated a novel central circuitry that interfaces between pain and itch and potentially integrates physiological information to modulate pain and itch through RVM neurons.

Conventionally, PBN has been identified as a sensory relay nucleus, which channels aversive/noxious sensations from the periphery to the forebrain regions to mount appropriate behaviors to prevent further harm and initiate coping behaviors to heal the injury. Recent studies have shown that activation of the PBN^Tacr1^ neurons, which receive direct inputs from the spinal cord Tacr1 expressing projection neurons, is sufficient to promote nocifensive behaviors and suppress itch (Deng et al., 2020; Barik et al., 2021). The itch suppression by PBN^Tacr1^ neurons could be due to their synaptic connections with limbic areas such as the hypothalamus, amygdala, or central thalamic nuclei (Barik et al., 2021; Pauli et al., 2022). However, our data indicate that the suppression primarily occurs through intra-brainstem projections to the RVM (Fig. 5). Notably, we found that the PBN^Tacr1^ downstream neurons in the RVM express the Tacr1 receptor (Fig. 3). Thus, a multi-synaptic neural circuit between the spinal cord, PBN, and RVM is instrumental in driving and modulating pain thresholds and simultaneously suppressing itch-induced scratching. The PBN^Tacr1^ neurons project to the central thalamus and may mediate the affective-motivational components of itch through this connection (Li et al., 2023).

Our fiber-photometry recordings indicate that the RVM^Tacr1^ neurons are tuned to noxious thermal stimuli, and the increased activity coincides with the licks and shakes (Fig. 6D-G). However, chemogenetic activation of the same cells was analgesic. This apparent inconsistency between the neural activity and behavioral outcome may be due to two reasons: 1) the RVM^Tacr1^ neurons are multimodal, and the fiber photometry technique is unable to resolve the diversity in these neurons 2) we found that the RVM^Tacr1^ neurons project to several brain regions including ones that drive pain-modulatory brain states such as anxiety, fear, and hunger (Rhudy and Meagher, 2000; Butler and Finn, 2009; Bushnell et al., 2013; Jennings et al., 2014; Alhadeff et al., 2018), thus compounding the behavioral phenotypes derived from RVM^Tacr1^ activation. In experiments underway, our efforts are focused on delineating the projection-specific roles of the RVM^Tacr1^ neurons in pain and itch modulation.

The Grp-Grpr, neuropeptide-GPCR expressing neurons and the molecules, play an essential role in facilitating and transmitting pruritus in the spinal cord (Sun and Chen, 2007; Sun et al., 2009). We found that Grpr is expressed in the lateral PBN and partially overlaps with Tacr1 neurons (Fig. 1A). Activating the PBN^Grpr^ neurons phenocopied artificial PBN^Tacr1^ activation in suppressing itch (Fig. 1D&G), and thus, the PBN^Grpr^ neurons may play an opposing role in modulating itch in the LPBN compared to the spinal Grpr neurons. In addition to the neuronal cell types in the PBN that express Grpr and Tacr1, the role of the neuropeptide ligands for these receptors Grp and Tac1, and the molecular signaling cascade induced by Grpr and Tacr1 activation in the PBN and RVM in pain and itch processing remain to be studied. The advent of fluorescence sensors enabling the study of neuropeptides (Wang et al., 2023) in live behaving mice can allow us to determine how and when neuropeptides Substance P (product of *tac1* gene) and Grp are released where the Tacr1 and Grpr are expressed in PBN and RVM. AAV-mediated delivery of silencing shRNAs (Liu et al., 2017; Rohn et al., 2024) targeting Tacr1 and Grpr can reveal the potential roles of these GPCRs in pain and itch. Notably, based on the results, the Tacr1 receptor and Tacr1-expressing neurons in the PBN and RVM can be targeted to modulate pain and itch.

## Methods

### Mouse lines

Animal care and experimental procedures were performed following protocols approved by the CPSCEA at the Indian Institute of Science. The animals were housed at the Central Animal Facility under standard transgenic animal housing conditions in a 12-hour light-dark cycle. Tacr1^Cre^ mice (Tacr1-T2A-Cre-Neo) were kindly donated by Dr. Hongkui Zheng, Allen Brain Institute. The Grpr^iCre^ mice were donated by Dr. Zhou-Feng Chen, Washington University, St. Louis. Genotyping for the mentioned strains was performed according to the protocols provided by Jackson Laboratories and the strain donors. All mice used in the behavioral assays were between 8 and 12 weeks old. All the behaviors were done during the light cycle.

### Viral Vectors

Vector used and sources: pAAV5-hsyn-DIO-EGFP (Addgene, Catalog# 50457-AAV 1), pAAV5-FLEX-tdTomato (Addgene, Catalog# 28306-AAV1), pAAV5-hsyn-DIO-hM3D(Gq)-mCherry (Addgene, Catalog# v141469), pAAV5-hsyn-DIO-hM4D(GI)-mCherry (Addgene, Catalog# 44362-AAV5), AAV9.syn.flex.GcaMP6s (Addgene, Catalog# pNM V3872TI-R(7.5)), v651-1), AAV9-DIO-PSD95-TagRFP (Donated by Mark Hoon, NIH), AAV5-hSyn-DIO-mSyp1_EGFP(University of Zurich, Catalog# v484-9), rAAV5-EF1α-DIO-oRVG (BrainVTA, Catalog# PT-0023), rAAV5-EF1α-DIO-EGFP-T2A-TVA (BrainVTA, Catalog# PT-0062), and RV-EnvA-Delta G-dsRed (BrainVTA, Catalog# R01002).

### Stereotaxic injections

Mice were anesthetized with 2% isoflurane/oxygen before and during the surgery. An incision was made to expose the skull, and subsequently, the skull was aligned to a horizontal plane. Craniotomy was performed at the marked point using a hand-held micro-drill (RWD). A Hamilton syringe (10 ul) with a glass pulled needle was used to infuse 300 nL of viral particles (1:1 in saline) at a rate of 100 nL/min. The following coordinates were used to introduce the virus: LPBN-AP: -5.34, ML: ±1.00, DV: -3.15; RVM-AP: -5.80, ML: +0.10, DV: -5.20. For rabies tracing experiments, rAAV5-EF1α-DIO-oRVG and rAAV5-EF1α-DIO-EGFP-T2A-TVA were injected first, followed by RV-EnvA-Delta G-dsRed after 3 weeks. Tissue was harvested after 1 week of rabies injection for histochemical analysis. The tissues in which obvious cell death through morphological examination was observed, were not included in the analysis. Post-hoc histological examination of each injected mouse was used to confirm that viral-mediated expression was restricted to the target nuclei.

### Fiber Implantation

Fiber optic cannula from RWD (Ø1.25 mm Ceramic Ferrule, 200 μm Core, 0.22 NA, L = 7 mm) were implanted at AP: -5.80, ML: +0.10, DV: -5.50 in the RVM after AAV-DIO-GCamp6s, was infused. Animals were allowed to recover for at least 3 weeks before performing behavioral tests. Successful labeling and fiber implantation were confirmed post hoc by staining for GFP for viral expression and injury caused by the fiber, respectively. Only animals with viral-mediated gene expression and fiber implantations at the intended locations, as observed in posthoc tests, were included in the analysis.

### Fiber Photometry

A dual-channel fiber photometry system from RWD (R810 model) was used to collect the data. The light from two light LEDs (410 and 470 nm) was passed through a fiber optic cable coupled to the cannula implanted in the mouse. Fluorescence emission was acquired through the same fiber optic cable onto a CMOS camera through a dichroic filter. The photometry data was analysed using the RWD photometry software and .csv files were generated. The start and end of stimuli were timestamped. All trace graphs were plotted from .csv files using GraphPad Prism software version 8.

### Behavioural Assays

#### Chloroquine-induced Itch Assay

The nape of the neck of mice was shaved 2-3 days before behavioral experimentations and the mice were habituated in the behavior room. Mice were individually placed in four-part plexiglass chambers with chamber dimensions of 6cm X 6cm X 14cm. The roof of the chamber had a hole for proper air ventilation. Animals were habituated in the chamber for 15 minutes before chloroquine injections. DCZ 2µg/kg body weight were administered intraperitoneally (i.p.) 15 min before chloroquine injection (Nagai et al., 2020). Chloroquine (375 µg/75 µl) was administered intradermally into the nape of the neck of the mice (Mishra and Hoon, 2013), and the subsequent scratching behavior was recorded for 30 minutes. Hind leg-directed scratching of the nape was characterized as a scratch.

#### Hotplate test

The thermal hotplate experiments were performed using the Hot and Cold Plate analgesiometer (HC-01, Orchid Scientific). The specifications of the instrument used are-enclosure size: 205 × 205 × 250 mm; plate size: 190 × 190 × 06 mm; temperature range: −5°C to 60 °C. A single experimenter introduced the mice in the enclosure on the thermal plate across all the experiments. The mice were habituated in the experimental room for 30 minutes and in the enclosure for 5 minutes at 32°C before the experimentation for three consecutive days. On the experimental day, mice were placed on the hotplate at 52°C temperature and the behavior were recorded for one minute using three Logitech web cameras, placed at the left, right, and front angles around the hotplate. Later, videos were quantified individually for any nocifensive behaviors (licks, shakes, and jumps) exhibited by the mice.

### Immunostaining, multiplex in situ hybridization, and confocal microscopy

Mice were anesthetized with isoflurane and perfused transcardially with 1X Phosphate Buffered Saline (PBS) (Takara) and 4% Paraformaldehyde (PFA) (Ted Pella, Inc.), consecutively for immunostaining experiments. Tissue sections were rinsed in 1X PBS and incubated in the blocking buffer (5% Bovine Serum Albumin (BSA) + 0.5% Triton X-100 + 1X PBS) for 1 hour at room temperature. Sections were then incubated in the primary antibody (dilution 1:1000 X in blocking buffer) at room temperature overnight. Sections were rinsed 1-2 times with 1X PBS and incubated for 2 hours in Alexa Fluor conjugated goat anti-rabbit/ chicken or donkey anti-goat/rabbit secondary antibodies along with DAPI (dilution 1:1000 X in blocking buffer) (Invitrogen) at room temperature, washed in 1X PBS, and mounted onto charged glass slides (Globe Scientific Inc.). Vecta Mount permanent mounting media (Vector Laboratories Inc.) was used for cover slipping the slides. Subsequently, sections were imaged on the upright fluorescence microscope (Khush Enterprises, Bengaluru) (2X, 4X, and 10X lens) and ImageJ/FIJI image processing software was used to process the images.

Fresh brains were harvested for in situ hybridization experiments. Multiplex ISH was done with a manual RNAscope assay (Advanced Cell Diagnostics). Probes were ordered from the ACD online catalogue. For the anatomical studies, the images were collected with 10X and 20X objectives on a laser scanning confocal system (Leica SP8 Falcon), and processed using the Leica image analysis suite software.

### Quantification and Statistical Analysis

All statistical analyses were performed using GraphPad PRISM 8.0.2 software. Student t-test and Two-way ANOVA tests were performed wherever applicable. ns > 0.05, ∗ P ≤ 0.05, ∗∗ P ≤ 0.01, ∗∗∗ P ≤ 0.001, ∗∗∗∗ P ≤ 0.0005.

## Funding

This research was funded by IndiaAlliance Intermediate Fellowship granted to A.B.

## List of Abbreviations

AAV: Adeno-associated virus
AHC: Anterior Hypothalamic Area
AITC: Allyl isothiocyanate
CeL: Central amygdala nucleus lateral division
CeM: Central amygdala nucleus medial division
CGRP: Calcitonin gene-related peptide
CM: Central medial thalamus
DCZ: deschloroclozapine
DM: Dorsal medial hypothalamus
DREADD: Designer Receptors Exclusively Activated by Designer Drugs
GFP: Green fluorescence protein
Gi: Gigantocellular reticular nucleus
Grp: Gastrin-releasing peptide
GRPR: Gastrin-releasing peptide receptor
IRt: Intermediate reticular nucleus
LH: Lateral hypothalamus
LHA: Lateral Hypothalamic area
LPBN: lateral parabrachial nucleus
LPO: Lateral Preoptic Area
MdD: Medullary reticular nucleus dorsal
MdV: Medullary reticular nucleus ventral
MPA: Medial preoptic area
mRt: mesencephalic reticular formation
MS: Medial Septum
p1RT: p1 reticular formation
PAG: Periaqueductal gray
PBN: Parabrachial nucleus
PCRt: Parvicell reticular nucleus
PF: Parafascicular thalamus
PLH: peduncular part lateral hypothalamus
PO: Preoptic area
RVM: rostral ventromedial medulla
SC: Superior Colliculus
SCP: superior cerebellar peduncles
SNR: Substantia Nigra
SoIV: solitary nucleus ventral
Tacr1: Tackykining receptor 1
VDB: nucleus of the vertical limb of the diagonal band
VGlut2: vesicular glutamate transporter 2
VLH: ventrolateral hypothalamic nucleus
VTA: Ventral Tegmental area

## Notes

### Competing Interest Statement

The authors have declared no competing interest.

